# Profiling virus-specific Tcf1+ T cell repertoires during acute and chronic viral infection

**DOI:** 10.1101/2020.03.20.000646

**Authors:** Alexander Yermanos, Ioana Sandu, Alessandro Pedrioli, Mariana Borsa, Franziska Wagen, Nathalie Oetiker, Suzanne P.M. Welten, Katharina Pallmer, Sai Reddy, Annette Oxenius

## Abstract

CD8 T cells play a crucial role in providing protection from viral infections. It has recently been established that a subset of CD8 T cells expressing Tcf1 are responsible for sustaining exhausted T cells during chronic lymphocytic choriomeningitis virus (LCMV) infection. Many of these studies, however, have been performed using T cell receptor (TCR) transgenic mice, in which CD8 T cells express a monoclonal TCR specific for the LCMV glycoprotein. To investigate whether the Tcf1+ and Tcf1-repertoires are naturally composed of similar or different clones in wild-type mice exposed to acute or chronic LCMV infection, we performed TCR repertoire sequencing of virus-specific CD8 T cells, including Tcf1+ and Tcf1-populations. Our analysis revealed that the Tcf1+ TCR repertoire is maintained at an equal or higher degree of clonal diversity despite harboring fewer cells. Additionally, within the same animal, there was extensive clonal overlap between the Tcf1+ and Tcf1-repertoires in both chronic and acute LCMV infection. We could further detect these virus-specific clones in longitudinal blood samples earlier in the infection. With respect to common repertoire parameters (clonal overlap, germline gene usage, and clonal expansion), we found minor differences between the virus-specific TCR repertoire of acute and chronic LCMV infection 40 days post infection. Overall, our results indicate that the Tcf1+ population emerging during chronic LCMV infection is not clonally distinct from the Tcf1-population, supporting the notion that the Tcf1+ pool is indeed a fuel for the more exhausted Tcf1-population within the heterogenous repertoire of LCMV-specific CD8 T cells.

## Introduction

Viral infections represent a global health problem and often invoke a potent immune response that can potentially result in severe immunopathology to the host if not counter-balanced by immunomodulatory circuits (Frebel *et al.*, 2012, 2010). Persistent viral infections can be differentiated as either actively replicating (hepatitis B virus, LCMV) or latent/reactivating infections (human immunodeficiency virus, hepatitis C virus, cytomegalovirus). Actively replicating chronic viral infections are characterized by a continuous high-level of viral antigen, which leads to excessive activation of virus-specific CD8 T cells. This potent and prolonged activation typically results in T cell exhaustion, where the upregulation of many inhibitory molecules concurs with a decrease in effector functions (Wherry, 2011; Frebel *et al.*, 2010). This phenomenon has been extensively researched in the LCMV murine infection model, in which both CD8 T cell-dependent clearance of acute (resolved within 2 weeks) and humoral mediated control of chronic (resolved after months) infection can be induced, depending on the dose and strain of the inoculating virus (Kräutler *et al.*, 2020). Following chronic LCMV infection, upregulation of inhibitory molecules such as PD-1, Lag3, Tim3 and loss of effector functions such as interferon gamma (IFN-y) and tumor necrosis factor (TNF) production occurs, impeding the CD8 T cell-mediated control of viral infection. Concomitantly, the adaptive host immune system emphasizes on the humoral response, characterized by pronounced differentiation (Osokine *et al.*, 2014; Fahey *et al.*, 2011) and sustained activity of T follicular helper cells, which are crucial for the eventual development of infection-resolving LCMV-neutralizing antibodies (Greczmiel *et al.*, 2017).

A population of CD8 T cells expressing the transcription factor T cell factor 1 (Tcf1) has been recently demonstrated to play an important role in the immune response during chronic LCMV infection. Absence of this population of Tcf1+ CD8 T cells resulted in reduced viral control and a decrease in virus-specific T cell clonal expansion (Utzschneider *et al.*, 2016). Current models suggest that this Tcf1+ population sustains the terminally differentiated, exhausted CD8 T cell pool during chronic LCMV infection, suggesting that there should be congruence between the polyclonal TCR repertoires of the Tcf1+ and Tcf1-populations. In addition to a role in chronic viral infection, Tcf1 has been implicated in the formation and function of effector CD8 T cells in acute LCMV infection (Wang *et al.*, 2019). Specifically, lack of Tcf1 resulted in an accelerated expansion and increased production of effector molecules such as IFN-y (Tiemessen *et al.*, 2014). Other studies have similarly demonstrated a role of Tcf1 in the formation of memory in the context of acute LCMV infection, where the clonal expansion of Tcf1 deficient CD8 T cells was impaired upon re-challenge and these Tcf1-cells were progressively lost over time (Jeannet *et al.*, 2010; Zhou *et al.*, 2010). It has been further demonstrated that checkpoint blockade results in the reinvigoration of the CD8 T cell response by effectively acting upon the Tcf1+ population in the context of both chronic viral infections and cancer (Utzschneider *et al.*, 2016; Siddiqui *et al.*, 2019), further stressing the importance of this T cell subset.

While multiple studies have recently investigated the transcriptional landscape of defined virus-specific Tcf1+ CD8 T cells in both acute and chronic infections (Utzschneider *et al.*, 2016; Chen *et al.*, 2019, 1), less attention has been devoted to studying the Tcf1+ and Tcf1-T cell receptor (TCR) repertoires under these two infection conditions. Advances in deep sequencing technologies enable the investigation of TCR (and antibody) repertoires at high-throughput and relatively inexpensive costs (Greiff, Miho, *et al.*, 2015; Georgiou *et al.*, 2014; Miho *et al.*, 2018). It is now feasible to profile the TCR repertoires overtime and across multiple individuals to reconstruct how host immunity adapts to foreign pathogens. It remains unknown, for example, how the TCR repertoire across multiple organs within the same host changes over time during acute and chronic viral infection – and whether and how repertoires diverge between functional different populations such as Tcf1+ and Tcf1-T cells. We therefore performed longitudinal sequencing of the LCMV-specific TCR repertoire following either chronic or acute LCMV infection, this included stratification into Tcf1+ or Tcf1-LCMV-specific T cells. Our analysis revealed that the Tcf1+ virus-specific CD8 T cell population maintained equal or greater diversity despite significantly fewer cell numbers than the corresponding Tcf1-population. We observed high congruence between the Tcf1+ and Tcf1-populations in both chronic and acute LCMV infection, demonstrated by parameters such as V gene usage, CDR3 length, clonal overlap, and clonal expansion. We furthermore demonstrated that many of the clonally expanded, virus-specific T cells found in spleen and lungs were detectable in the blood at various sampling time points during the infection. Taken together, our results both describe the TCR repertoire following acute and chronic infection and support the hypothesis that the Tcf1+ population sustains the Tcf1-repertoire.

## Results

### Quantitative and qualitative differences in Tcf1 populations following acute and chronic LCMV infection

Previous research demonstrating phenotypic differences in the polyclonal T cell population of Tcf1+ and Tcf1-subsets during acute or chronic infection lead us to question whether these subsets share similar or distinct TCR repertoires. To address this, we performed longitudinal sequencing on the TCR variable beta chain (TRB) repertoire in nine mice with acute or chronic LCMV infection; mice were of the transgenic Tcf1 reporter strain (*Tcf7*^GFP^, which expresses GFP under the control of the *Tcf7* promoter that encodes Tcf1 (Utzschneider *et al.*, 2016)) (Figure 1A). This included both unsorted T cells from serial blood sampling and LCMV-specific T cells from lungs and pooled spleen and lymph nodes (spleen/LN), which were isolated by fluorescently activated cell sorting (FACS) based on specificity towards the immunodominant peptide gp^33-41^. In both chronic and acute LCMV infection, gp^33-41^-specific cells were sub-sorted into Tcf1+ and Tcf1-populations, in combination with PD-1 expression (in the case of chronic LCMV infection) and CD44 expression (in case of both acute and chronic infection) (Figures 1B, 1C). Specifically, those PD-1^hi^ CD8 T cells lacking specificity towards the gp^33-41^ epitope (GP33-) were sorted from mice with chronic LCMV infection to include cells with other LCMV-specificities. While this sorting strategy resulted in a discrepancy between infection cohorts for those T cell populations unspecific to gp^33-41^, it nevertheless allowed the comparison between Tcf1+ and Tcf1-repertoires across repertoires skewed towards LCMV-specificity. Overall, our experimental pipeline resulted in 16 distinct T cell populations (8 of which with known LCMV-specificity to the gp^33-41^ epitope) for TCR repertoire sequencing ranging from hundreds to hundreds of thousands of cells (Figures 1D, 1E). FACS analysis demonstrated lower frequencies and reduced total numbers of Tcf1+ CD8 T cells in both the spleen/LN and lungs of chronically infected mice relative to the acute infection (40 dpi) (Figures 1F, 1D, 1E). As expected given the role of Tcf1 in orchestrating a central memory-like phenotype (Zhou *et al.*, 2010), the pooled spleen/LN samples for both infection cohorts had a higher proportion of Tcf1+ than the lungs (Figure 1F). Further analysis confirmed that the majority of GP33-specific CD8 T cells were PD-1^hi^ in both spleen/LN and lungs from chronically infected mice, whereas hardly any GP33+ T cells expressed PD-1 in acutely infected mice 40 dpi (Figures 1G, 1H). FACS analysis confirmed phenotypic differences between the two infection cohorts and thus we proceeded with TCR repertoire sequencing to quantitatively characterize the underlying clonal diversity of the Tcf1+ and Tcf1-T cell populations.

**Figure 1.**
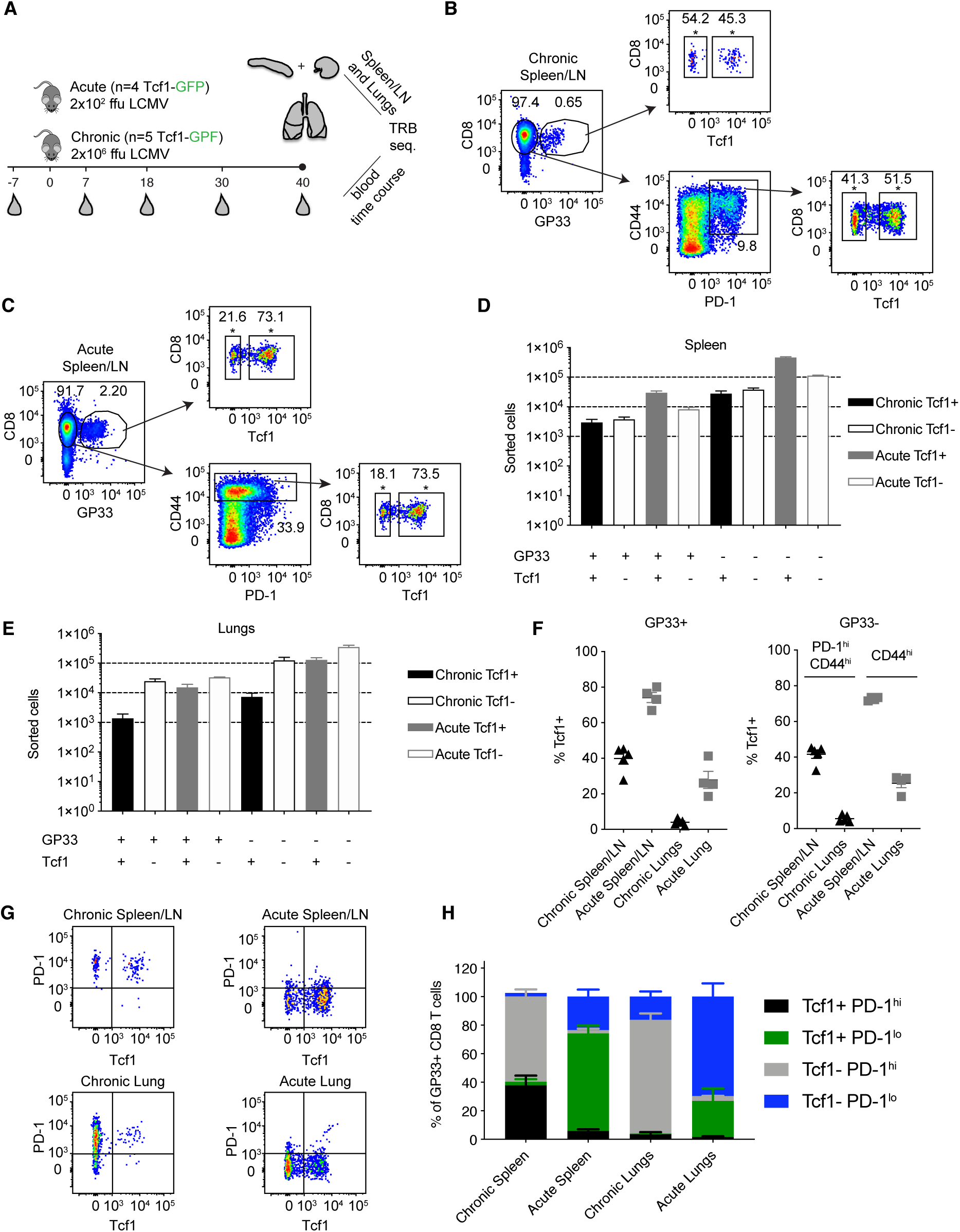
Experimental setup for TCR-beta repertoire sequencing. (A) Both unsorted blood samples and sorted T cells from pooled spleen/LNs and lungs from acutely (2×10^2^ focus forming units (ffu) and chronically (2×10^6^ ffu) infected *Tcf7*^GFP^ mice were sampled for TCR-beta repertoire sequencing. (B-C) Representative FACS plots for sorting strategy for chronic and acute LCMV infection cohorts. Asterisks (*) indicate the final population for sequencing. (D-E) Sorted cell numbers for each of the populations included in TCR-beta sequencing experiments. Black bars indicate chronically infected mice and gray bars indicate acutely infected mice. Error bars indicate standard error of the mean. n=4-5 mice for acute and chronic cohorts, respectively. (F) Frequency of Tcf1+ CD8 T cells in the pooled spleen/LN or lungs of chronically and acutely infected mice 40 dpi. Error bars indicate standard error of the mean. n=4-5 mice per experiment. (G) Representative FACS plots displaying the relationship between Tcf1+ and PD-1 of GP33+ CD8 T cells 40 dpi. (H) Quantification of G. Error bars indicate standard error of the mean. n=4-5 mice per experiment.

### Increased CD8 clonal diversity in Tcf1+ fraction in chronic compared to acute LCMV infection

Recent work has demonstrated that the Tcf1+ CD8 T cell population sustains the Tcf1-population during chronic infection (Utzschneider *et al.*, 2016). We therefore predicted the clonal diversity of the Tcf1+ TCR repertoire to be at least as diverse as the Tcf1-repertoire. Enumerating the total number of unique clones (defined by identical CDR3b amino acid sequence) revealed that the Tcf1+ population contained at least equal or greater number of unique clones than the Tcf1-population across all organs and for both acute and chronic infection (Figures 2A, 2B). This was true whether looking at the clonal diversity within the immunodominant GP33+ epitope or in those repertoires of antigen-experienced cells where TCR specificity was unknown (GP33-). To investigate the sensitivity of our sequencing pipeline to varying levels of input material, we next compared the total number of clones extracted from the sorted CD8 T cell populations to the blood samples, in which no cell sorting was performed. We expected the blood T cell repertoires of uninfected mice (containing both CD8 and CD4, sampled 7 days before infection) to contain much higher levels of clonal diversity than that observed in the spleen/LN and lung repertoires. Indeed, this analysis revealed that the blood time points from the same sequencing batch contained on average over 20,000 unique clones compared to the thousands of sorted cells from the spleen and lungs (Figure 2C).

**Figure 2.**
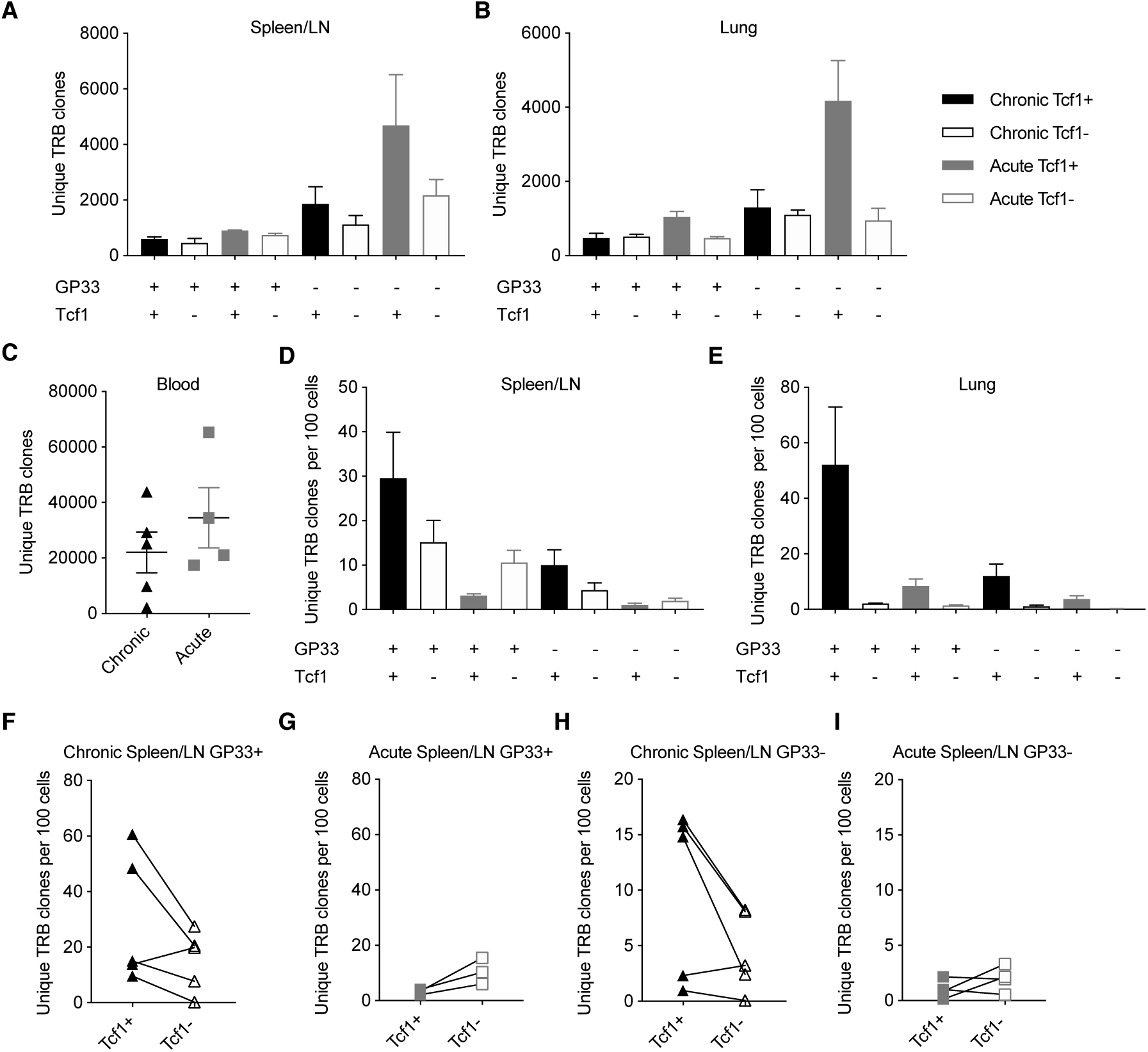
Increased or equal clonal diversity in Tcf1+ repertoire compared to Tcf1-in chronic but not acute infection. (A-B) Number of unique TRB clones in the spleen/LN and lung samples sorted 40 days post infection (dpi). Clone is defined as unique CDR3b amino acid sequence. (C) Number of unique TRB clones in the blood 7 days before infection. (D-E) Unique TRB clones normalized by the number of sorted cells 40 dpi. Error bars indicate standard error of the mean. (F-I) Unique TRB clones normalized by the number of sorted cells 40 dpi for the Tcf1+ and Tcf1- repertoires for each mouse. n=4-5 mice per experiment.

Our initial analysis suggested that the highest degree of clonal diversity in GP33+ TCR repertoires was in the lungs (Figures 2A, 2B). This analysis had initially disregarded, however, that these repertoires arose from varying cell numbers (Figures 1D, 1E). We therefore questioned whether the Tcf1+ repertoires contained more unique clones than their corresponding Tcf1-counterparts even after normalizing for sorted cell number. While the spleens of the chronically infected cohort tended to have more unique clones per sorted cells in the Tcf1+ compared to the Tcf1-population, this was reversed in the acute infection in both GP33+ and GP33-repertoires (Figure 2D). Performing the same analysis in the lungs, however, revealed that Tcf1+ repertoires for both acute and chronic infections contained more unique clones than their corresponding Tcf1-repertoires in both GP33 positive and negative populations. While not as robust as in the lung repertoires, the trend that the Tcf1+ repertoire had a higher number of unique clones after normalizing by cell number was observed in the majority of spleens when looking within individual mice for both GP33+ and GP33-populations, further suggesting that clonal diversity is maintained in the Tcf1+ population (Figures 2F-I).

### Comparable germline gene usage in the Tcf1+ and Tcf1-repertoires

After observing the comparable clonal diversity in the Tcf1+ and Tcf1-TCR repertoires, we hypothesized that these repertoires would use similar germline V-gene elements. We therefore quantified the percentage of unique clones using a particular V-gene in the virus-specific Tcf1+ and Tcf1-spleen/LN populations in chronically infected mice (Figure 3A). Both repertoires had the highest number of unique clones with TRBV29, which composed approximately 20% of the unique TRB clones. Many of the lowly expressed TRB V-genes (e.g. TRBV3, TRBV26, TRBV24, TRBV23) were similarly absent in both Tcf1+ and Tcf1-repertoires, suggesting that there were no dramatic differences in recruitment and selection of the two virus-specific T cell populations. We next quantified the mean V-gene usage across all animals for each of the sorted cell populations, which demonstrated considerable similarity across the various TCR repertoires (Figures 3C, 3D). Indeed, certain V-genes such as TRBV1 and TRBV29 were dominant in the majority of spleen and lung repertoires regardless of the population in chronically infected animals (Figures 3C, 3D). We next questioned whether quantifying this distribution of V-genes would reveal subtle but discernable differences between the repertories. To address this, we calculated the Shannon evenness, a metric commonly used to describe the distribution of clonal frequencies (Greiff, Bhat, *et al.*, 2015). This analysis similarly demonstrated comparable frequencies of V-genes for the virus-specific splenic repertoires of both acute and chronic infection (Figure 3E).

**Figure 3.**
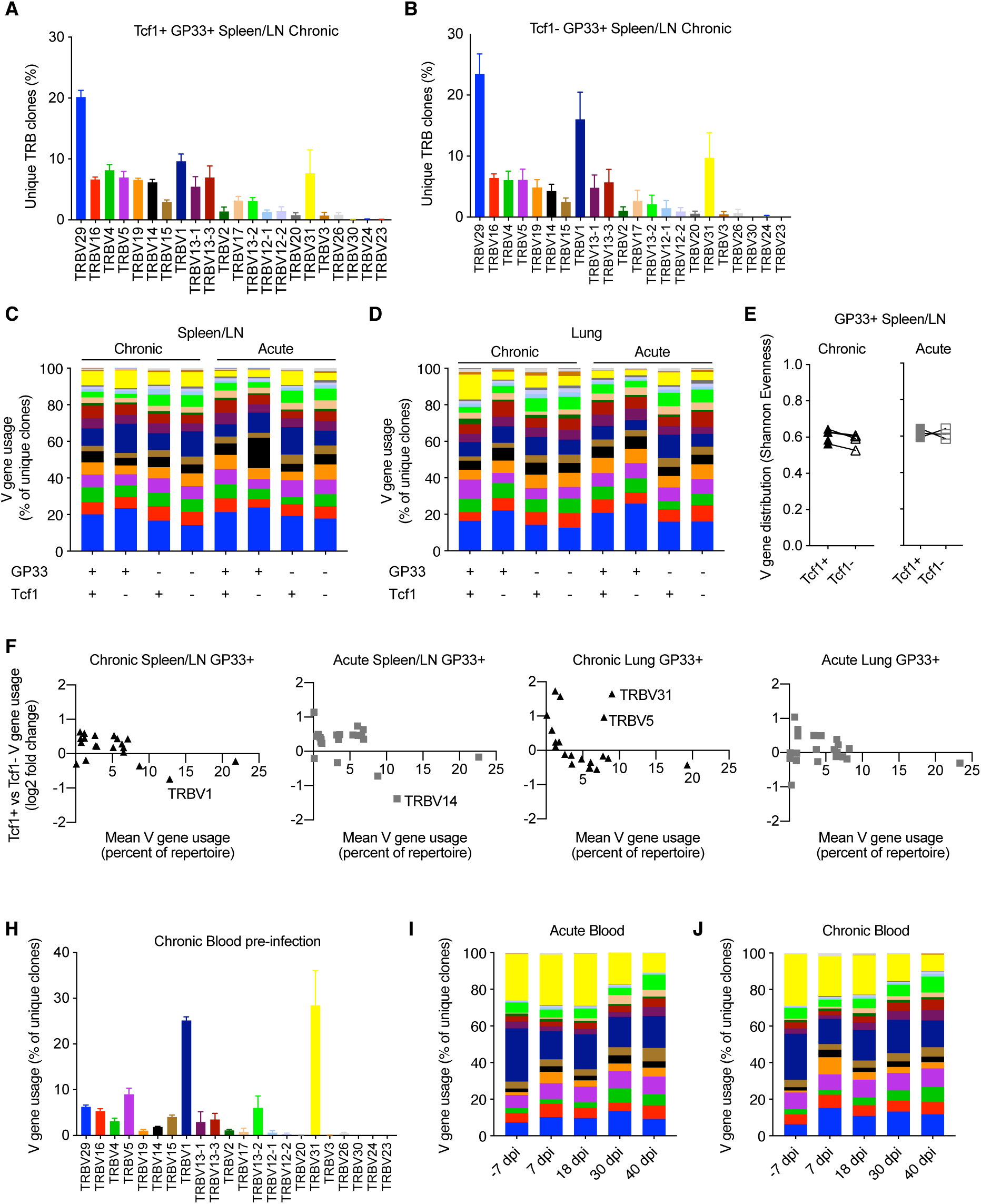
Similar germline gene usage of Tcf1+ and Tcf1- repertoires. (A-B) Example TRB V-gene usage for the Tcf1+ and Tcf1- virus-specific repertoires from the spleen/LN 40 days post infection (dpi). Error bars indicate standard error of the mean. n=4-5 mice per experiment. (C-D) Mean V-gene usage in each sequenced cohort determined by GP33 specificity and Tcf1 expression. (E) Shannon evenness quantifying the distribution of V-gene usage in the splenic repertoires. Values closer to 1 indicate even distributions of V-gene usage, whereas values closer to 0 indicate highly variable distributions of V-gene usage. (F) Up- and down-regulation of particular V-genes in Tcf1+ versus Tcf1- negative repertoires. Each point represents a different TRB V-gene segment. Y values greater than 0 indicate that a given V-gene is on average upregulated in the Tcf1+ repertoire relative to the Tcf1- repertoire. (H) Example V-gene usage from bulk blood repertoires from the five mice in the chronic LCMV cohort before any viral infection. (I-J) Average V-gene usage in the blood repertoires at each time point for acute and chronic infection. n=4-5 mice per experiment.

We next explicitly quantified whether certain TRB V-genes were more or less present in either the Tcf1+ or Tcf1-repertoires for either infection cohort. To relate the expression level to V-gene (up- or down-regulation), we plotted the mean V-gene usage against the log_2_ fold-change of V-gene usage between the Tcf1+ and Tcf1-within the same organ and infection cohort (Figure 3F). Overall, the differences in the V-gene usage of virus-specific clones was comparable across all groups, with very few highly employed V-genes (e.g. more than 5% of the repertoire) with a log_2_ fold-change greater than 1/-1. While there were two V-genes up-regulated in the lungs (TRV31 and TRBV5) in the Tcf1+ compared to the Tcf1-repertoires of chronically infected mice, this upregulation was not similarly observed in the spleen (Figures 3F, S2A). Similarly, TRBV14 was more abundant in the Tcf1-repertoires in the spleens of acutely infected mice compared to the Tcf1+ counterparts, despite this trend being absent in the corresponding lung repertoires (Figures 3F, S2A).

Surprised by the high degree of similarity in TRB V-gene usage across all of our samples, we questioned whether similar patterns of germline gene usage would be detected in the blood, particularly prior to infection. Quantifying the percent of unique clones aligning to each V-gene revealed a substantial (>40% of unique clones) portion of the unsorted blood repertoire consisted of TRBV1 and TRBV31 before viral infection. While these two genes were present in the sorted populations 40 dpi, neither consisted of over 20% of the repertoire on average as seen in the blood repertoires of both infection cohorts before any viral infection (Figures 3A-D, 3H, S2A, S2B). We hypothesized that the blood repertoire V-gene usage would become more similar to the distribution observed at the terminal time points. At the later blood time points (30 and 40 dpi), both cohorts demonstrated a decrease in genes TRBV31 and TRBV1 (Figures 3I, 3J), reaching comparable levels to those observed in the lungs and spleen 40 dpi (Figures 3C and 3D). We finally determined whether such congruence between infection cohorts and Tcf1+ and Tcf1-repertoires was also observable when looking at other repertoire parameters, such as the length of complementarity determining region 3 (CDR3). Indeed, our analysis revealed highly similar distributions of CDR3b lengths for both splenic and lung repertoires between the Tcf1+ and Tcf1-repertoires (Figure S3A). In contrast to the V-gene usage, the blood CDR3 length distribution was highly similar to those found at terminal time points following LCMV infection (Figure S3B), suggesting that LCMV infection did not selectively recruit naïve clones with a particular CDR3 length. Quantifying the length distribution of the GP33-repertoires revealed minor differences between the difference Tcf1+ and Tcf1-populations. Taken together, these data demonstrate convergent repertoire fingerprints across both Tcf1+ and Tcf1-subsets for both cohorts, consistent with the hypothesis that the pool Tcf1-CD8 T cells are sustained by the Tcf1+ population.

### Mouse-specific clones in Tcf1+ and Tcf1-populations

After observing the consistencies involving V-gene usage and CDR3 length between the Tcf1+ and Tcf1-repertoires, we next quantified the extent to which the two repertoires shared common clones (within or between individual mice). We hypothesized that the two repertoires would share a high proportion of clones given the hypothesis that the Tcf1+ pool sustains the Tcf1-population. We first calculated the percent of Tcf1-clones found in the Tcf1+ repertoires of either the same mouse or other mice within the same cohort (Figures 4A, 4B). This analysis revealed considerable numbers of public clones within the organ-specific repertoires between individual mice within either infection cohorts. Despite the high clonal overlap across different mice, we observed a mouse-specific trend in which the Tcf1-repertoire had more clones in common with the Tcf1+ repertoire of the same mouse for both spleen and lungs (Figures 4A, 4B). Comparing this to the number of clones shared in the bulk blood repertoires across individual mice within the same infection cohort revealed that, on average, there was a smaller fraction of cohort-shared blood clones compared to the terminal, virus-specific repertoire (Figures 4A, 4B, 4C).

**Figure 4.**
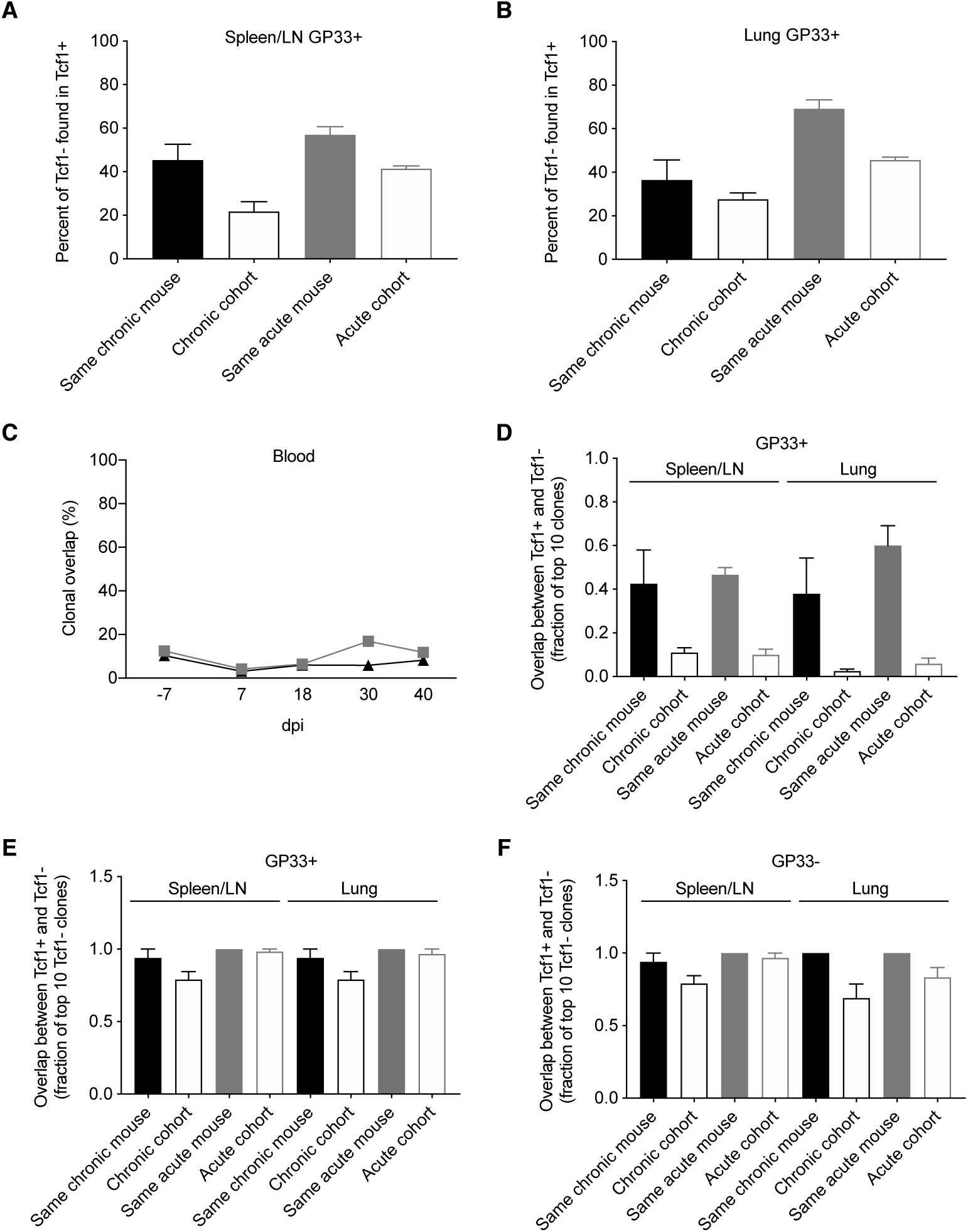
Clonal overlap between the Tcf1+ and Tcf1- repertoires. (A-B) Percent of the GP33+ Tcf1- repertoire found in the GP33+ Tcf1+ repertoire from either the same mouse or other mice in the same infection cohort for spleen/LN and lung repertoires. Clonal overlap was defined by identical CDR3b amino acid sequences found in both repertoires. (C) Clonal overlap between bulk blood repertoires at the various time points throughout the infection. (D) Clonal overlap of the top expanded clones in the Tcf1+ and Tcf1- virus-specific repertoires. The fraction of the top 10 most expanded GP33+ clones in the Tcf1+ repertoire that were shared between the top 10 most expanded GP33+ clones in the Tcf1- repertoire of either the same mouse or between two mice in the same cohort. (E) The fraction of the top 10 most expanded GP33+ clones in the Tcf1- repertoire that were found at any clonal frequency in the GP33+ Tcf1+ repertoire of either the same mouse or between two mice in the same cohort. (F) The fraction of the top 10 most expanded GP33- clones in the Tcf1- repertoire that were found at any clonal frequency in the GP33- Tcf1+ repertoire of either the same mouse or between two mice in the same cohort.

Despite the aforementioned mouse-specific overlap between the Tcf1- and Tcf1+ populations, we were surprised that only ∼50% of the clones were common to both repertoires. This prompted us to question whether restricting our analysis on the most expanded clones of both Tcf1+ and Tcf1-repertoires would increase the clonal overlap between these populations. We thereby quantified the fraction of the top 10 most expanded clones in either the Tcf1+ and Tcf1-repertoires. This analysis demonstrated a much stronger trend of mouse-specific overlap, in that approximately 4-6 of the top 10 GP33+ clones were common across the Tcf1+ and Tcf1-repertoires of the same mouse, but only approximately 0-1 of the top 10 clones were common to different mice within the same cohort (Figure 4D), suggesting that expanded clones in the Tcf1+ repertoire feed into the Tcf1-repertoire within an individual mouse. We next questioned how relaxing the restriction to the top 10 clones of the Tcf1+ repertoire would impact this clonal overlap. We thereby quantified the fraction of the top 10 expanded GP33+ clones from the Tcf1-fraction found in the entire Tcf1+ repertoire, regardless of clonal expansion. We observed that the majority of the 10 most expanded clones were found at some degree of clonal expansion in the Tcf1+ repertoire (Figure 4E), again further strengthening the hypothesis that the Tcf1+ repertoire sustains the Tcf1-repertoire. Surprisingly, the majority of these highly expanded, virus-specific clones were shared across other mice within the same infection cohort at either comparable or slightly less frequencies (Figure 4E), which may represent a commonly generated TCR sequence that could be maintained in both Tcf1+ and Tcf1-repertoires based on affinity. We finally questioned whether this was similarly true for the GP33-populations for both acute and chronic infection and observed results consistent with the GP33+ repertoires (Figure 4F). Taken together, the higher mouse-specific overlap between Tcf1+ and Tcf1-repertoires, particularly in the top 10 most expanded virus-specific clones, further suggests an equilibrium between the two compartments within a single animal.

### Expanded clones found across multiple organs and time points

After observing a high degree of clonal overlap between the Tcf1+ and Tcf1-repertoires within the same organ, we next determined the extent of clonal overlap between the spleen/LN, lungs and blood repertoires. We hypothesized that that there should be a high overlap between the Tcf1-lung repertoire and the Tcf1+ central memory pool, as it has been shown that these Tcf1+ virus-specific T cells reside primarily in secondary lymphoid organs (Utzschneider *et al.*, 2016; Im *et al.*, 2016). To answer this, we quantified clonal overlap using the Jaccard index, a metric which takes into account a different number of clones, and ranges between 0 and 1, where 1 indicates complete overlap and 0 represents no overlap. While there was a trend that the lung Tcf1- and the spleen/LN Tcf1+ repertoires contained the highest degree of overlap compared to other Tcf1+ vs Tcf1-permutations, the Jaccard indices were comparable across groups (Figure 5A). Furthermore, there was a surprising fraction of shared clones across animals within the same cohort (Figure 5A). Performing a similar analysis for the acutely infected animals revealed increased overlap between the spleen/LN and lung repertoires compared to chronically infected mice (Figure 5A). We next questioned if restricting our analysis to the top 10 most expanded clones between the spleen/LN and lungs would inform whether they were present in both the central lymphoid and lung T cell populations. Indeed, we observed a strong mouse-specific trend in which the most expanded clones were more often shared between central and peripheral organs in the same mouse relative to other mice from the infection cohort (Figure 5B). There was again a minor trend of increased overlap between the lung Tcf1- and the spleen/LN Tcf1+ repertoires relative to other Tcf1+/- combinations, particularly compared to the overlap between the lung Tcf1+ and spleen Tcf1-population (Figure 5B), further supporting the Tcf1+ population as a source of the Tcf1-repertoire.

**Figure 5.**
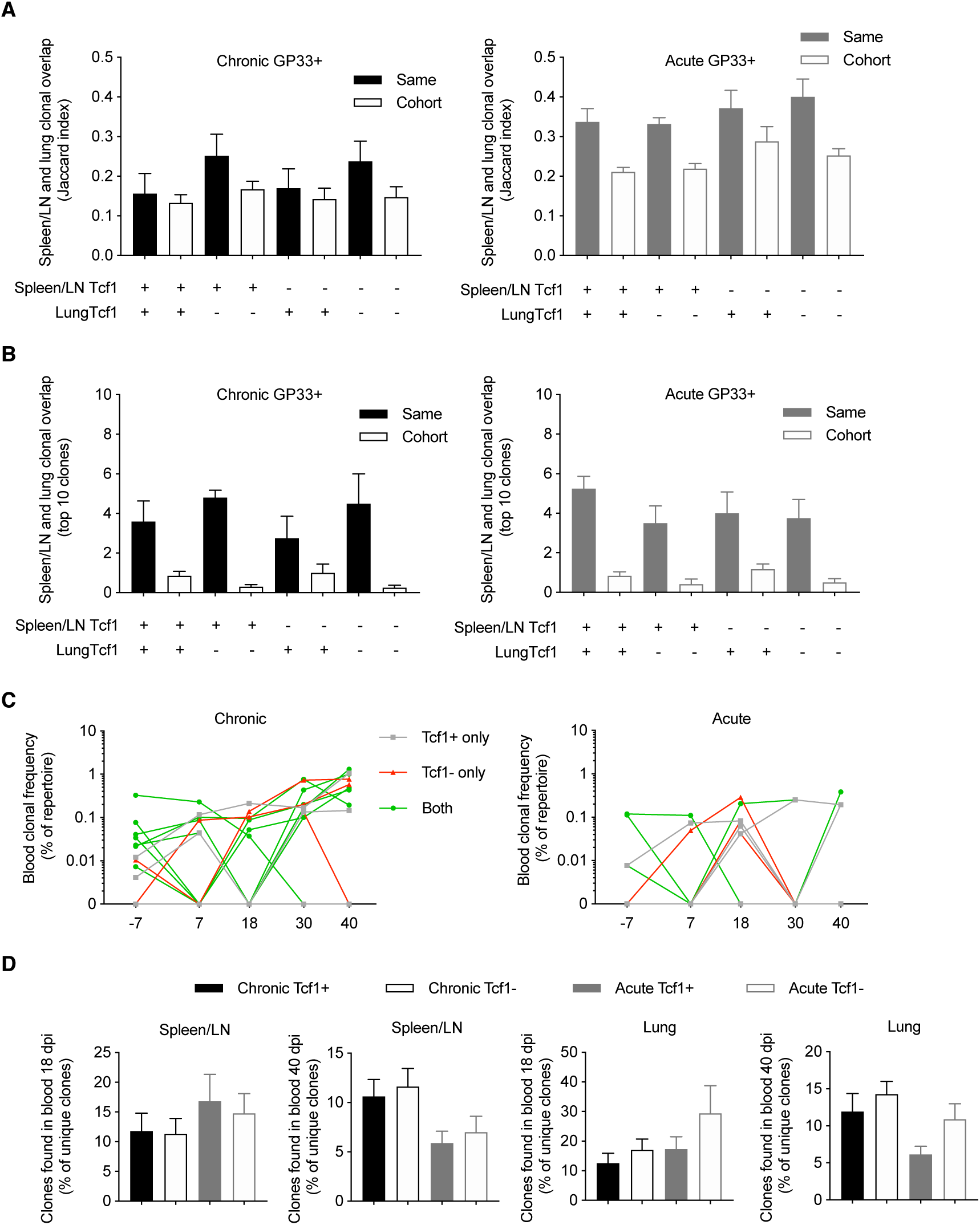
Clonally expanded virus-specific T cells detected in the spleen/LN, lungs, and blood repertoires. (A) Jaccard index quantifying the clonal overlap between the spleen/LN and lung repertoires of either the same animal (filled color) or other animals within the same cohort (no color fill). Higher Jaccard indices indicate higher clonal overlap (defined by unique CDR3b amino acid sequences). Key below graph refers to whether Tcf1+ or Tcf1- of the spleen/LN and lungs were compared. (B) Clonal overlap between the top 10 most expanded clones in the Tcf1+ and the top 10 most expanded clones in the Tcf1- repertoires between spleen/LN and lungs of either the same animal (filled color) or other animals within the same cohort (no color fill). (C) The clonal frequencies of two example mice (% of total blood repertoire) of the top 10 most expanded clones from the spleen GP33+ repertoire. Color indicates if the clone was shared between Tcf1+ and Tcf1- repertoires within the same mouse. (D) Clonal overlap between the blood repertoires 18 and 40 dpi and terminal sorted populations of the same animals. Error bars indicate standard error of the mean. n=4-5 mice per experiment.

We subsequently investigated whether the most expanded clones found at the terminal time point could be detected in the blood TCR repertoires of the same mice for both chronic and acute infection cohorts. Indeed, our time-resolved sequencing showed that many of top 10 most expanded clones from both Tcf1+ and Tcf1- of spleen/LN repertoires were observed at earlier time points (Figures 5C, 5D). Given blood sampling captures only a fraction of the total circulating T cells, it is somewhat expected there were several lowly expressed clones not found at each time point (Figure 5C). We were nevertheless curious as to whether there was a difference in clonal overlap between acute and chronic infection given early (approximately two weeks) viral clearance in acute but not chronic infection. We therefore quantified the overlap between our virus-specific sorted T cell population either 18 or 40 dpi, expecting higher overlap between those clones found 40 dpi in the blood and terminal organs in chronic infection compared to acute infection. Indeed, there was a trend that chronically infected mice had a higher percent of clones shared between both the blood repertoires 40 dpi and the spleen/LN and lung repertoires also collected 40 dpi (Figure 5D). Taken in concert, these results further demonstrate mouse-specific overlap between the most expanded clones found in both the spleen/LN and lung repertoires, and a large fraction of both Tcf1+ and Tcf1-clones were present in the blood repertoire.

### Minor differences in shared clones, V-gene usage, and clonal expansion between chronic and acute LCMV infection

The majority of our analysis until now has focused on the differences between the Tcf1+ and Tcf1-repertoires, regardless of infection cohort. However, the observation that TCR repertoires were more similar to the blood 40 dpi in chronically infected mice (Figure 5D) lead us to question whether additional differences were detectable between the two infection cohorts. We first quantified the clonal overlap between the two infection conditions and compared the fraction of shared clones (identical CDR3b amino acid sequence) to the Jaccard index of each cohort individually (Figure 6A). In the spleen for both the GP33+ and GP33-repertoires, there was comparable overlap between cohort-restricted (chronic vs chronic, acute vs acute) and mixed-cohorts (acute vs chronic) in both Tcf1+ and Tcf1-repertoires (Figure 6A). Of note is that the Jaccard index was on average higher for GP33+ repertoires in both organs, indicating that clones specific for the immunodominant epitope are more likely to be found across mice than clones potentially unspecific or targeting a different viral epitope (Figure 6A). Performing the same quantification for the GP33+ repertoires in the lungs revealed there were very few clones (approximately 3% and 7% for Tcf1+ and Tf1-, respectively) shared between the two cohorts (Figure 6A). Similarly, the mixed-cohort overlap was on average lower than the cohort-restricted overlap for the GP33-lung, albeit with all samples demonstrating lower Jaccard indices than the GP33+ population.

**Figure 6.**
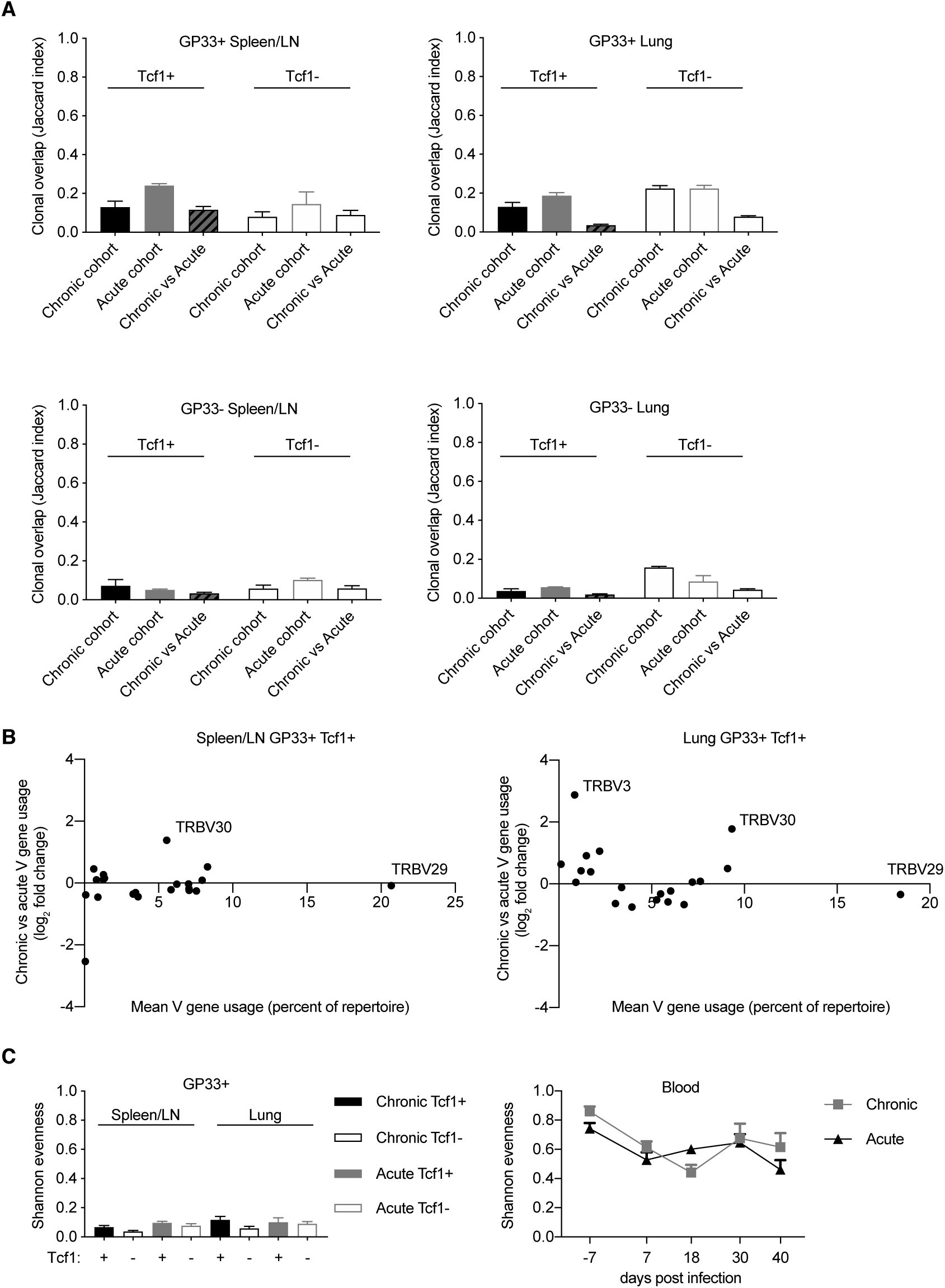
Minor differences between T cell repertoires from acute and chronic infection cohorts. (A) Jaccard index quantifying the clonal overlap between the repertoires of either animals within the same infection cohort (chronic versus chronic, acute versus acute) or across infection cohorts (chronic versus acute). (B) Up- and down-regulation of particular V genes in chronic versus acute GP33+ Tcf1+ repertoires. Each point represents a different TRB V gene segment. Y values greater than 0 indicate that a given V gene is on average upregulated in the chronic GP33+ Tcf1+ repertoires relative to the acute GP33+ Tcf1+ repertoires. (C) Shannon evenness quantifying clonal expansion in the GP33+ and blood repertoires. Higher values indicate more similar levels of clonal expansion across the repertoire.

After observing organ-specific differences between the mixed-cohort clonal overlap, we questioned whether this lack of clonal overlap corresponded to different germline gene usage. We therefore plotted the V-gene usage (mean percent of repertoire) versus the log_2_ fold-change of V-gene expression between chronic and acute infection. We had expected more pronounced differences in V-gene usage in the lung relative to the spleen/LN repertoires based on the aforementioned clonal overlap analysis (Figure 6A). There were minor differences between the V-gene usage of the two infection cohorts in both spleen/LN and lungs, and in fact, we even noticed a single V-gene, TRBV30, that was upregulated in chronic infection relative to acute infection in both spleen/LN and lung repertoires (Figure 6B). This and our previous clonal overlap analyses (Figures 6A, 6B) focused primarily upon unique clones, thereby ignoring potential effects of clonal expansion. However, after observing a mouse-specific increase in clonal overlap in the most expanded clones across various analyses (Figures 4, 5), we determined whether any metrics quantifying clonal expansion could differentiate chronic and acute infections in either the spleen/LN, lungs or blood repertoires. To this end, we calculated the Shannon evenness describing the distribution of clonal expansion for all clones in each repertoire (Greiff, Bhat, *et al.*, 2015). For all sorted cell populations, there were large variations in clonal expansion (represented by Shannon evenness indices close to 0), suggesting that some clones had undergone extensive replication and others were largely unexpanded (Figure 6C). There was a trend of higher clonal expansion in the GP33+ Tcf1+ compared to the Tcf1-repertoires, particularly in chronic infection (Figure 6C). Indeed, comparing this to the Shannon evenness from the blood repertoires revealed that these terminal sorted cell populations were much more expanded than any of the blood time points (Figure 6C). There was a decrease in the Shannon evenness following viral infection for both cohorts, presumably due to the expansion of virus-reactive cells. While we observed a trend of increased expansion in the chronically infected cohort relative to those acutely infected animals 40 dpi, there was considerable variability throughout the duration of the sampling, which could be partially explained by biological sampling depth. In conclusion, these analyses further demonstrate minor differences between chronic and acute infection TCR repertoires while simultaneously revealing further differences between organ-specific and Tcf1+/- TCR repertoires.

## Discussion

While normally expressed in naïve and memory CD8 T cells, Tcf1 has been recently demonstrated to distinguish a memory-like population of T cells which play a role in resolving both chronic infection and cancer due to their ability to feed into the pools of more differentiated /exhausted CD8 T cells and due to their ability to primarily respond to check-point blockade. In the case of chronic LCMV infection, initial clonal expansion of virus-specific T cells in Tcf1 knock-out mice were comparable to wild type mice but were unable to resolve viral infection (Utzschneider *et al.*, 2016). Despite certain memory-like characteristics of Tcf1+ CD8 T cells, such as re-expansion potential, low cytotoxicity, and expression of cytokines such as IL-2, these cells paradoxically co-express markers typically characterized by an “exhausted” phenotype, such as Lag-3 and PD-1, in addition to global gene transcriptional signatures (Utzschneider *et al.*, 2016). Most relevant to the experiments presented in this study, it was shown that the Tcf1+ population aid both viral and tumor control by sustaining the production of effector Tcf1-cells (Siddiqui *et al.*, 2019; Utzschneider *et al.*, 2016). These previous findings suggest a high degree of similarity between the Tcf1+ and Tcf1-repertoires, particularly following chronic LCMV infection. Our experimental approach enabled us to quantify several repertoire metrics, including the extent of clonal overlap, clonal diversity and germline gene usage. Furthermore, leveraging deep sequencing we were able to compare TCR repertoires following acute and chronic infection of CD8 T cells isolated from different organs.

Overall, our findings support a model in which the Tcf1+ population sustains the Tcf1-repertoire during chronic LCMV infection, particularly demonstrated by an equal or greater number of unique clones in the Tcf1+ population (especially after normalizing for cell numbers) and that almost all of the most expanded Tcf1-clones were found in the corresponding Tcf1+ counterpart. These findings were further supported by similar V-gene usage, CDR3 length and the extent of clonal expansion between Tcf1+ and Tcf1-repertoires. Furthermore, there was a trend in chronically infected mice of increased clonal expansion in the Tcf1+ repertoire relative to the Tcf1-counterpart (Figure 6C), again consistent with previous reports demonstrating higher population expansion in the Tcf1+ compartment on a monoclonal level (Utzschneider *et al.*, 2016). Our analysis additionally revealed a large fraction of virus-specific clones that were shared across mice, which were present at much higher frequencies than observed in the antibody repertoire of B cells in a similar study (Kräutler *et al.*, 2020). This may be somewhat expected given that T cells rely on a preformed TCR repertoire that cannot mutationally evolve during antigenic challenges. This is in contrast to the B cell response, which relies upon antibody sequence diversification through somatic hypermutations such that repertoires dramatically diverge between hosts to eventually attain “personalized” neutralization capacities after months of infection (Staupe *et al.*, 2019; Kräutler *et al.*, 2020).

In contrast to the antibody repertoires following LCMV infection, we observed minor differences in the TCR repertoires between acute and chronic cohorts, including the upregulation of the TRBV30 in both spleen/LN and lung repertoires in chronic infection (Figure 6B) and virus-specific clones restricted to a particular infection cohort (Figure 6A). Overall, the repertoire structure was strikingly similar between the two infection conditions, further suggesting that phenotypic diversity plays a larger role than CD8 T cell repertoire diversity in response to LCMV infection (Chen *et al.*, 2019; Siddiqui *et al.*, 2019; Hudson *et al.*, 2019). Further experiments investigating the CD4 T cell landscape may provide more apparent differences between the two infection cohorts and Tcf1+ and Tcf1-populations, given both the role of T follicular helper cells in chronic infection and that previous studies performing CD4 depletion have demonstrated that the terminally differentiated Tcf1-population shrinks but not in the proliferation-competent CD8 T cells (Greczmiel *et al.*, 2017; Kanev *et al.*, 2019). Despite previously reported differences in clonal expansion kinetics of virus specific CD8 T cells in regards to chronic vs acute LCMV infection (Wherry *et al.*, 2003), we saw comparable degrees of clonal expansion in both blood repertoires when incorporating clonal information (Figure 6C). Clonal expansion metrics from bulk repertoire sequencing experiments should be interpreted with caution, however, given it is difficult to determine if the sequence-derived clonal frequency actually correlates to cell number (Horns *et al.*, 2020; Khan *et al.*, 2016). We have therefore based the majority of our analysis on unique TRB CDR3 sequences, without any information regarding the TCR variable alpha (TRA) chain. TCRb sequencing studies are better suited to compare repertoires between conditions rather than comment on absolute values of unique clones and clonal expansion due to amplification biases during PCR and sequencing errors (Khan *et al.*, 2016). Despite these shortcomings, we observed hundreds of unique GP33+ clones, which is in line with previous studies that have quantified comparable numbers of GP33+ naïve precursors in the entire CD8 repertoires of uninfected mice (Jenkins and Moon, 2012; Kotturi *et al.*, 2008; Obar *et al.*, 2008). This suggests that the majority of these naïve precursor cells are recruited and maintained during LCMV infection. However, the use of technologies better suited to absolute quantification of sequence diversity and clonal expansion would be helpful to validate these among other findings. For example, employing single-cell sequencing platforms, where it is possible to pair the transcriptional landscape with paired TRB and TRA sequences while maintaining error correction through the use of unique molecular identifiers (Horns *et al.*, 2020; Singh *et al.*, 2019), would better resolve the relationship between clonal expansion, gene expression, and repertoire diversity.

## Methods

### Mice

All animal experiments were performed according to institutional guidelines and Swiss Federal regulations and were approved by the veterinary office of the canton of Zurich (animal experimentation permissions 115/2017). 18-20 week old female *Tcf7*^GFP^ mice (Utzschneider *et al.*, 2016) were infected with 200 ffu or 2×10^6^ ffu LCMV clone 13 i.v. to induce acute and chronic infections, respectively. All mice were housed under specific-pathogen-free conditions in individually ventilated cages and were not involved in experiments outside of this study. Bedding and nesting material were provided for enrichment. 4-5 mice were housed within the same cage for all experiments.

### LCMV virus production

LCMV clone 13 was produced as previously described (Kräutler *et al.*, 2020). In brief, LCMV clone 13 was propagated by infection of BHK-21 cells for 24-48h. Supernatant was filtered through 0.22 µm filter (TPP filtermax) and then resuspended in sterile PBS (GIBCO).

### Blood and organ isolation

200 µL of blood was sampled from the leg vein at indicated time points (excluding 7 dpi, in which 50 µL of blood was sampled) during the course of infection. Upon sacrifice 40 dpi, blood was sampled via intracardiac bleeding. Blood was collected in heparin-coated microtainers (BD) and centrifuged for 5 minutes at 2000 g to separate plasma and cells. Pellets were resuspended in 50 µL PBS and mixed with 1500 mL Trizol LS (Ambion) for future RNA extractions. Spleens and six lymph nodes (axial, inguinal, salivary gland) were pooled together. Mice were perfused with PBS to remove blood associated cells before extracting the lungs. Single cell suspensions from spleens and LN pools were prepared by meshing the tissue through a 70 µM cell strainer. Lungs were first cut into small pieces, incubated with Collegenase I and DNAse I for 30 minutes, meshed through 70 µM strainers then followed by a 30% percoll gradient. Cells were incubated with fluorescently conjugated antibodies for 30 minutes at 4°C. Antibodies used in the experiments; CD8-PerCP (BioLegend; Cat#100794), CD44-PE (BioLegend; Cat#103008), PD-1-PECy7 (BioLegend; Cat#135215). MHC Class 1 tetramers for gp^33-41^ was prepared and conjugated to APC as previously described (Altman *et al.*, 1996). Multi-parametric flow cytometric sorting was performed using FACSAria with FACSDiva software. Sorted cells were lysed in 600 µL Trizol LS (Ambion) and stored at -80°C until RNA-extraction. FACS data was analyzed using FlowJo software (Tree Star).

### Sequencing library preparation

RNA was extracted using the Direct-zol RNA MiniPrep kit (Zymo) according to manufacturer’s instructions without the use of DNAse. First strand cDNA was synthesized in a total volume of 20 µl using 11.5 µl of RNA, 0.5 µl oligo(dT) primers (100 mM, life technologies), 1 µl dNTPs (10 mM, life technologies), 1 µl 0.1M DTT (life technologies), 1 µl RNAsin Plus RNAse inhibitor (10K, Promega AG), 1 µl Superscript III (200 U/ml, life technologies) and 4 µl 5x Superscript III buffer for 10 min at 50°C, 10 min at 25°C and 60 min at 50°C. Polymerase was inactivated by incubation for 5 min at 94°C. TCR sequencing libraries were then prepared in a two-step PCR approach amplifying the TCR-β chain as previously described (Dash *et al.*, 2011) using 19 TRBV forward primers and 1 TRBC reverse primer. The first PCR was performed using Q5 Hotstart Polymerase HiFi (NEB) in a reaction volume of 25 µl with overhang-extended primers under the following conditions (5 × 65°C, 35 × 62°C). Indexed Illumina sequencing adaptors were added during a second PCR step (5 × 40°C, 25 × 65°C). Following magnetic bead clean up (CleanNGS), amplicons were eluted in 15 µl of buffer. The quality of the libraries was assessed using a fragment analyzer (Bioanalyzer, Agilent). Libraries were pooled and sequenced across three runs (run 1 contained all acute sorted samples with chronic mice 1 and 2; run 2 contained the remaining chronic samples plus blood samples before any infection; run 3 contained the remaining blood time points) on the Illumina MiSeq platform with 2 × 300 cycles. Due to technical problems one Tcf1+ GP33+ repertoire from an acutely infected mouse and one blood sample from a chronically infected mouse 7 dpi were not included in the analysis.

### TCR analysis

Paired-end sequencing fastq files were processed using the MiXCR software (v3.0.1) (Bolotin *et al.*, 2015) with reads aligned to the built-in murine reference genome as previously described (Kräutler *et al.*, 2020). Following error correction using default parameters, clonotyping was performed on identical nucleotide CDR3 regions using MiXCR. Clonotypes containing identical CDR3 amino acid sequences were subsequently merged into a final clonotype. Further filtering was performed to include only those in-frame TCRb clones with V and J gene alignments and that were supported by more than one sequencing read as performed previously (Greiff *et al.*, 2017). V-, D- and J-gene assignment was based on the germline segment with the highest alignment score, and in the case of ties the first gene was selected. Log_2_ fold-change ratios were calculated based on the mean proportion of repertoire for each V gene within a given infection cohort. When relating log_2_ fold-change to mean V-gene usage, mean V gene usage was calculated across all mice included in the comparison, regardless of cohort. Repertoire parameters such as Shannon evenness and Jaccard indices were calculated as previously described (Kräutler *et al.*, 2020; Greiff, Bhat, *et al.*, 2015). Briefly, Shannon evenness was calculated by first using diversity function from the R package vegan supplying the proportional clonal frequency for each repertoire (Oksanen *et al.*, 2019). We then took the exponential of the output and divided this by the total number of unique clones for each repertoire, as previously described (Greiff, Bhat, *et al.*, 2015). The Jaccard indices were calculated by dividing the clonal overlap between two repertoires by the total diversity shared between the two samples. Clonal overlap was exclusively defined as two clones sharing identical CDRH3 amino acid sequence.

