## Supplementary materials for "Profiling virus-specific Tcf1+ T cell repertoires during acute and chronic viral infection"

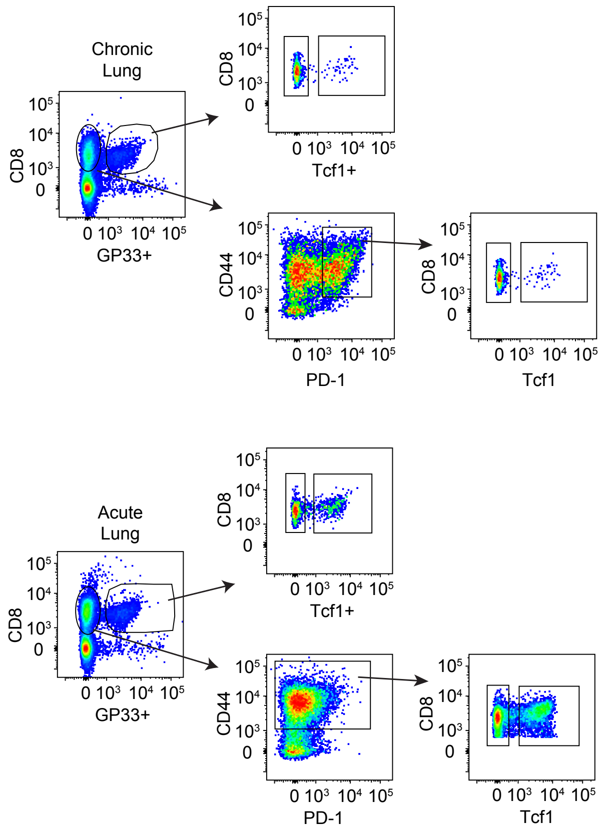


**Figure S1. Example sorting strategy and representative FACS plots for lungs of chronically and acutely LCMV infected *Tcf7*$^{GFP}$ mice.**

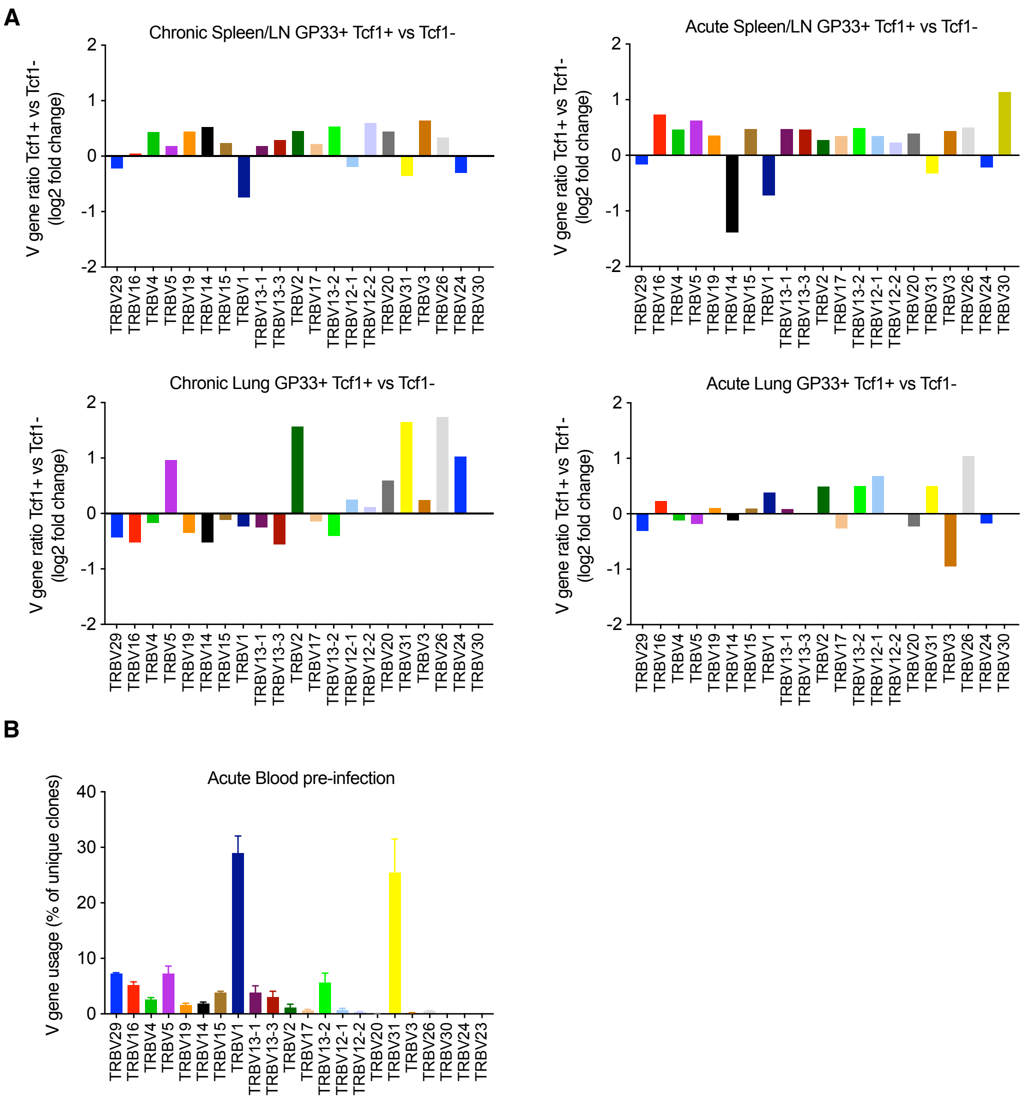


**Figure S2. Minor changes in V gene usage between Tcf1+ and Tcf1- repertoires.** (A) The $log_2$ change between the Tcf1+ and Tcf1- repertoires for each V gene from sorted spleen/LN and lungs 40 dpi. (B) V gene usage (percent of unique clones) for the acute blood repertoire before any viral infection.

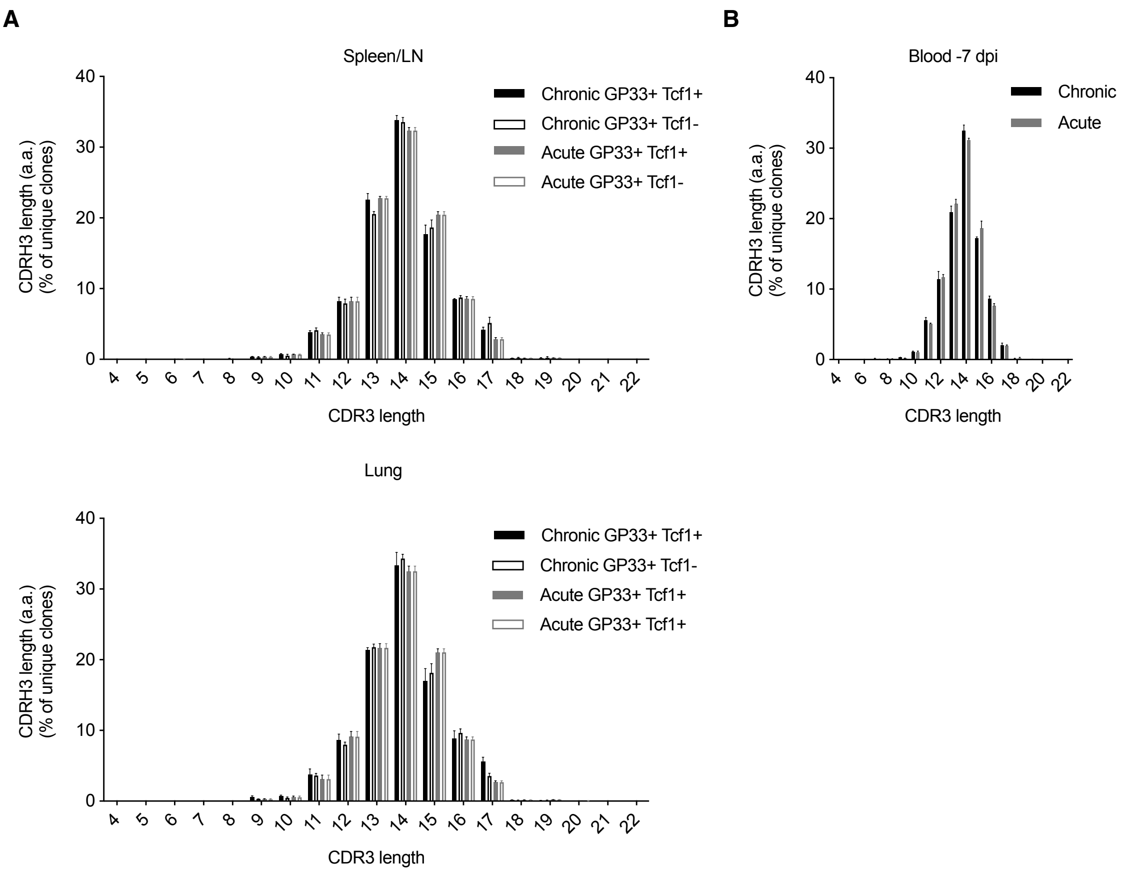


**Figure S3. Similar CDR3b length distribution across Tcf1+ and Tcf1- repertoires.** (A) CDR3b length distribution for (A) Spleen/LN and lung 40 days post infection (dpi) and (B) blood before any viral infection.
